# Circulating microRNA biomarkers for metastatic disease in neuroblastoma patients

**DOI:** 10.1101/309823

**Authors:** Fjoralba Zeka, Alan Van Goethem, Katrien Vanderheyden, Fleur Demuynck, Tim Lammens, Anneleen Decock, Hetty H Helsmoortel, Joëlle Vermeulen, Rosa Noguera, Ana P Berbegall, Valérie Combaret, Gudrun Schleiermacher, Geneviève Laureys, Alexander Schramm, Johannes H Schulte, Sven Rahmann, Julie Bienertová-Vašků, Pavel Mazánek, Marta Jeison, Shifra Ash, Michael D Hogarty, Mirthala Moreno-Smith, Eveline Barbieri, Jason Shohet, Frank Berthold, Frank Speleman, Matthias Fischer, Katleen De Preter, Pieter Mestdagh, Jo Vandesompele

**Affiliations:** Center for Medical Genetics, Ghent University, Ghent, Belgium; Cancer Research Institute Ghent (CRIG), Ghent University, Ghent, Belgium; Department of Pediatric Hematology-Oncology and Stem Cell Transplantation, Ghent University Hospital, Ghent, Belgium; Pédiatrie, Hôpital de Jolimont, Haine-Saint-Paul, Belgium; Department of Pathology, Medical School, University of Valencia/CIBERONC Madrid, Spain; INCLIVA Biomedical Research Institute, Valencia, Spain; Laboratoire de Recherche Translationnelle, Centre Léon-Bérard, Lyon, France; Department of Paediatric Oncology, Institut Curie, Paris, France; Molecular Oncology, West German Cancer Center, University Hospital Essen, University of Duisburg-Essen, Essen, Germany; Pediatric Oncology and Hematology, Charité University Medicine, Berlin, Germany. German Cancer Research Center (DKFZ), German Cancer Consortium (DKTK), Heidelberg, Germany; Genome Informatics, Institute of Human Genetics, University Hospital Essen, University of Duisburg-Essen, Essen, Germany; Department of Pediatric Oncology, University Hospital Brno, Brno, Czech Republic; Pediatric Hematology Oncology, Schneider Children’s Medical Center of Israel, Tel Aviv, Israel; Division of Oncology, The Children’s Hospital of Philadelphia, Philadelphia, United States; Department of Pediatrics, Section of Hematology-Oncology, Texas Children’s Cancer Center, Baylor, College of Medicine, Houston, Texas, United States; Pediatric Oncology and Hematology, University Children’s Hospital of Cologne, Medical Faculty, University of Cologne, Cologne, Germany; Center for Molecular Medicine Cologne (CMMC), University of Cologne, Cologne, Germany

## Abstract

In this study, the circulating miRNome from diagnostic neuroblastoma serum was assessed for identification of non-invasive biomarkers with potential in monitoring metastatic disease. After determining the circulating neuroblastoma miRNome, 743 miRNAs were screened in two independent cohorts of 131 and 54 patients. Evaluation of serum miRNA variance in a model testing for tumor stage, MYCN status, age at diagnosis and overall survival, revealed tumor stage as the most significant factor impacting miRNA abundance in neuroblastoma serum. Differential expression analysis between patients with metastatic and localized disease revealed 9 miRNAs strongly associated with metastatic stage 4 disease in both patient cohorts. Increasing levels of these miRNAs were also observed in serum from xenografted mice bearing human neuroblastoma tumors. Moreover, murine serum miRNA levels were strongly associated with tumor volume, suggesting this miRNA signature may be applied to monitor disease burden.

## Introduction

Neuroblastoma is a childhood cancer arising from the developing sympathetic nervous system.[1] The disease is responsible for 11% of pediatric cancer deaths and is ranked third among cancers with the highest mortality rate.[2] Approximately 60% of the neuroblastoma patients are diagnosed with metastatic disease, mostly represented in the high-risk patient sub-group. High-risk patients have low survival rates of about 50%, and two thirds of these children will ultimately relapse, eventually entering a second line of intensive treatment, enrolling in an early-stage clinical trial or receiving palliative therapy.[1] Monitoring of treatment response is essential for assessment of its efficacy and for guiding the procedure of follow-up treatment after refractory or relapsed neuroblastoma.

Current methods for monitoring disease and evaluation of treatment response are mostly based on detection of minimal residual disease (MRD), which is determined by estimating the number of residual tumor cells in peripheral blood or bone marrow.[3] While several cellular MRD markers have been proposed, biomarkers for non-invasive cell-free determination of disease burden are currently lacking.

MicroRNAs, a class of small non-coding RNAs involved in various aspects of health and disease, including neuroblastoma pathophysiology, seem to be particularly stable and abundant in serum and plasma.[4,5] Several studies have reported a relationship between serum miRNAs and disease characteristics such as *MYCN* amplification, or increased risk for adverse outcome.[6,7] The interest in pursuing the investigation of the miRNome from liquid biopsy samples is also fortified by the fact that collection, processing and storage of these samples is relatively straightforward and less invasive. For these reasons, the circulating miRNome may be a very valuable source for development of more advanced or refined tests for longitudinal treatment follow-up of neuroblastoma patients.

In the present study, we quantified the circulating miRNome from 185 diagnostic serum samples and assessed the association of four disease characteristics with serum miRNA abundance: the international neuroblastoma staging system (INSS), gene copy number status of the *MYCN* oncogene, age at diagnosis, and overall survival. We found a strong positive correlation between serum miRNA abundance and tumor stage and selected 9 miRNAs with a proportional increase with increasing tumor stage. As tumor stage reflects cancer spread, we propose this set of miRNAs as serum markers for assessment of disease burden in human neuroblastoma.

## Results

### Defining the human neuroblastoma circulating miRNome

As only a subset of miRNAs end up in circulation, we decided to first define the circulating miRNome (i.e. the ensemble of miRNAs that are detectable in serum from NB patients). To this end, we created three serum pools, each composed of 5 patient samples (Figure 1). Pooled samples were either high-risk patients that died of the disease (pool 1), high-risk patients that survived (pool 2) or low-risk patients that survived the disease (pool 3). We argue that these groups are representative for the majority of the neuroblastoma patient population, resulting in a comprehensive selection of circulating NB miRNAs. The pooled samples were screened for 1805 miRNAs. A set of 751 miRNAs, well expressed in at least one of the serum pools, was ultimately selected, hereby defining the NB miRNome. Expression of the NB miRNome was subsequently analyzed by RT-qPCR in 2 NB patient cohorts of 131 and 54 serum samples, respectively. Before normalization, eight miRNAs with missing values in more than 75% of the samples were excluded from the dataset. All analyses were performed on the first cohort initially. The second cohort was used to validate the findings from the first cohort.

**Figure 1:**
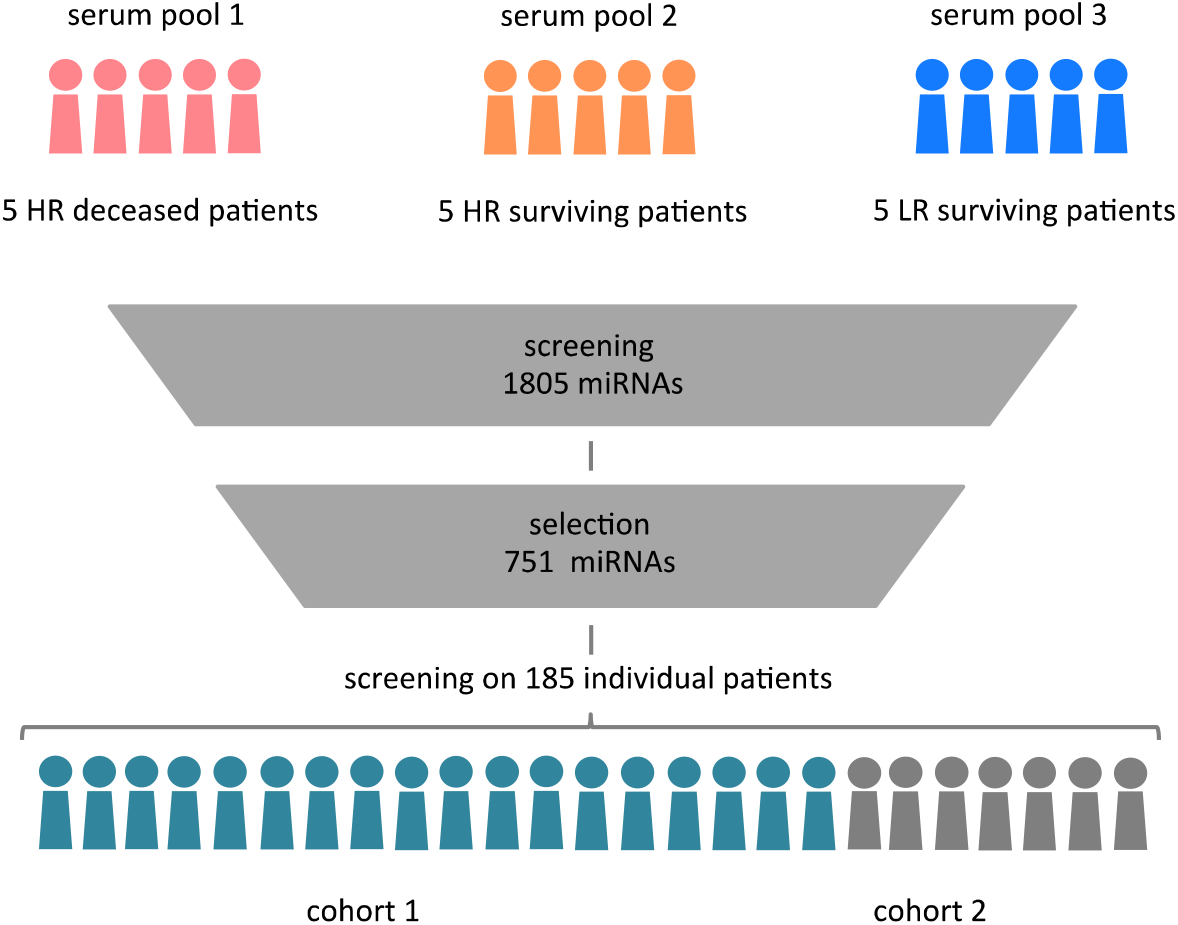
Three serum pools were prepared, each containing 5 serum samples from three neuroblastoma sub-groups: 5 high-risk (HR) deceased patients, 5 high-risk surviving patients and 5 low-risk (LR) surviving patients for serum pool 1, 2 and 3, respectively. Expression of 1805 miRNAs was measured by RT-qPCR. 751 well-expressed miRNAs were selected and profiled on two independently collected and processed patient cohorts of 131 patients and 54 patients, respectively.

### Metastatic disease status has the largest impact on circulating miRNA levels in serum

Generalized additive modeling (GAM) was used on normalized miRNA expression values in order to quantify the association with tumor stage, *MYCN* status, age at diagnosis and overall survival. A straightforward model was chosen, whereby interaction terms were excluded, since our aim was to determine which of these features has the largest impact on serum miRNA expression and hence is responsible for the largest variance in the data set.

The analysis returned the level of significance (P-value) and the percentage of variance in the dataset explained by each of the tested features. The level of significance was used to measure the prediction ability of disease features for the 743 tested miRNAs. The percentage of variance was used to define the importance of each feature in influencing miRNA expression in serum. The average statistical significance, the number of significant miRNAs and the average variance explained by each disease feature is provided in Figure 2.

**Figure 2:**
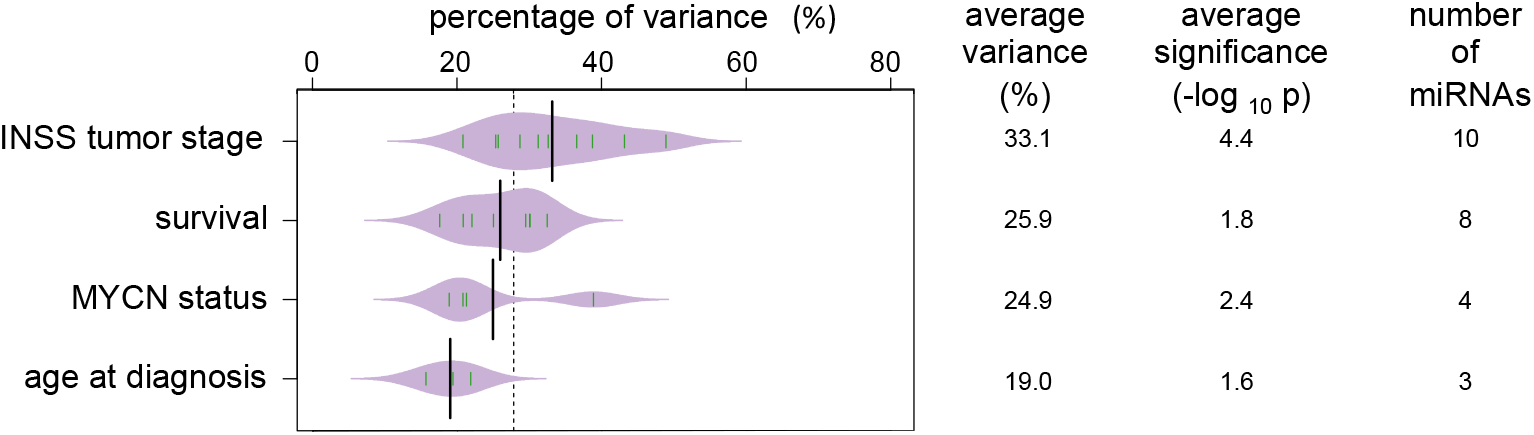
Bean plot of the average variance per disease feature (%), average significance (−log 10 of the P-value) and number of miRNAs significantly influenced by disease features. Green lines in the violin plot represent percentage of variance for individual miRNAs. The evaluated disease features were tumor stage (1,2, 3, 4 and 4S), overall survival (survival), *MYCN* status and age at diagnosis (18 moths cut-off). P-values were corrected for multiple-testing error by the Benjamini-Hochberg’s-method prior to averaging.

INSS tumor stage explains on average 33.1% of the variance in this serum miRNA dataset. Ten miRNAs were identified as being strongly significantly affected by tumor stage. The average level of significance was P-value < 0.0001 (range: 3.65 × 10^−2^ − 3.19 × 10^−11^). Overall survival status (alive or dead of disease) explained an average 25.9% of the variance. Eight miRNAs were identified with an average significance of P-value < 0.05 (range: 3.58 × 10^−2^ − 5.88 × 10^−3^). *MYCN* copy number status and age at diagnosis were responsible for 24.9% and 19.0% of the expression variance, respectively, with an average significance of P-value <0.01 and <0.05, respectively. The number of miRNAs to be significantly influenced by *MYCN* status or age at diagnosis were 4 and 3, respectively.

None of the identified miRNAs were significantly influenced by two or more disease characteristics. Based on the overall percentage of variance explained and the average level of significance for the identified sets of miRNAs for each disease feature, we concluded that INSS tumor stage had the largest impact on circulating serum miRNA levels.

### Identification of serum miRNAs as potential markers for disease burden

Mann-Whitney U test was used to select differentially expressed miRNAs between serum from patients with metastatic disease (stage 4 tumors) and serum from patients with localized disease (stage 1 and stage 2 tumors)

The analysis revealed 327 differentially expressed miRNAs (after correction for multiple-testing). 184 and 143 miRNAs were more abundant in serum from metastatic and localized disease, respectively (Figure 3a). The magnitude of the differences was much larger for miRNAs circulating in metastatic serum samples (Figure 3b). In fact, metastatic serum revealed 10 miRNAs with minimal fold-changes higher than 2 ΔCq units (~4-fold difference), while the maximum fold-change in serum from localized disease was lower than 2 ΔCq units.

**Figure 3:**
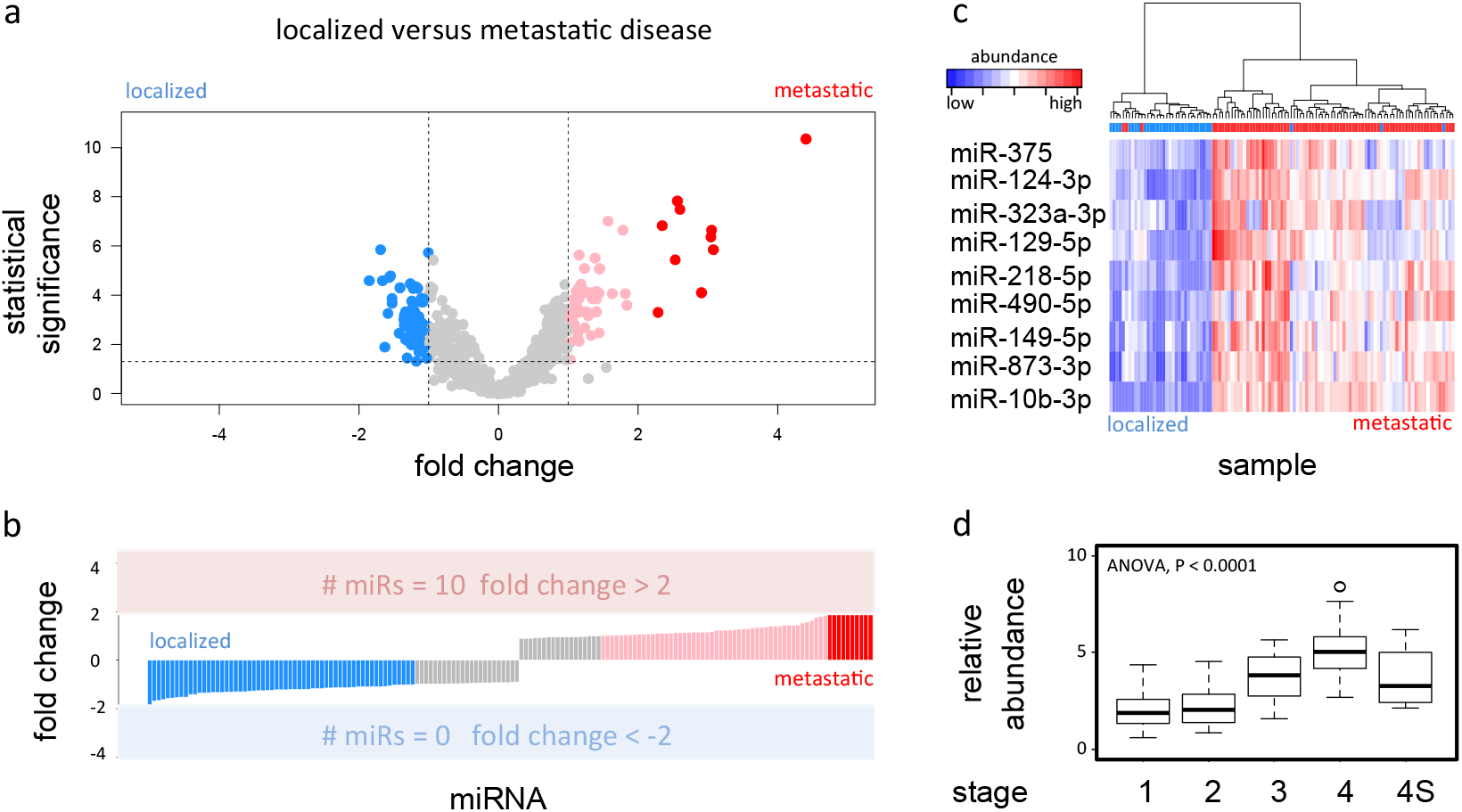
**a)** Fold change (log2 scale) of miRNA abundance in serum (x-axis) against statistical significance (y-axis, −log10 of the P-value) between patients with metastatic (stage 4) neuroblastoma (n=76) and patients with localized (stage 1 and stage 2) neuroblastoma (n=33). Higher fold change denotes higher abundance in metastatic serum. Pink dots represent miRNAs with at least two-fold (1 log2 unit) higher abundance and red dots miRNAs with at least a four-fold higher abundance in serum from metastatic patients; blue dots represent miRNAs with at least two-fold higher abundance in serum from patients with localized disease. For the selection of differentially abundant miRNAs, a Mann-Whitney U test was used at 0.05 significance level (corrected for multiple-testing by the Benjamini-Hochberg’s method). **b)** Fold change distribution of miRNA abundance in localized versus metastatic serum. The bar colors correspond to dot colors in plot (A). **c)** Heatmap depicting hierarchical clustering of serum samples (cohort 1) based on 9 miRNAs with at least four-fold higher abundance in serum from metastatic patients. **d)** Boxplot of average miRNA expression for the selected set of 9 miRNAs in serum from stage 1, 2, 3, 4 and 4S patients.

Interestingly, 9 out of the top 10 differential miRNAs were also put forward by the GAM analysis. These overlapping miRNAs were miR-873-3p, miR-149-5p, miR-124-3p, miR-218-5p, miR-490-5p, miR-323a-3p, miR-10b-3p, miR-375, and miR-129-5p (Figure 3c).

We further evaluated the serum levels of the 9 overlapping miRNAs in patients stratified according to INSS tumor stage. Average miRNA abundance was calculated per sample, samples were grouped per stage and depicted by a boxplot in Figure 3d.

A proportional increase of miRNA levels in serum from stage 1 to stage 4 patients was observed. Analysis of variance (Anova-test) was used to quantify the significance of the difference between stage groups. This analysis revealed a significant P-value of 2.0 × 10^−16^. All differences were significant except for the differences between stage 1 and 2 tumors, and stage 4S and all other tumor stages.

Even though we would label the selected miRNAs as being indicators of metastatic tumor load (stage 4 disease) there is a discrepancy with respect to how the miRNAs behave in serum from stage 4S patients. Stage 4S patients have a particular pattern of metastatic disease with good outcome due to spontaneous regression; yet the expression of the selected set of 9 miRNAs is not significantly different from the localized tumor stages. This may be due to the fact that this cohort only contained a limited number of samples. Alternatively, it may reflect the difference in biology between stage 4 and 4S tumors, which may result in a different miRNA signature in patient serum samples. Based on these results, we assume that the identified set of miRNAs qualify as candidate markers of metastatic, non-stage 4S disease (hereafter referred to as markers for metastatic disease).

### Validation of serum miRNA markers for metastatic disease in independent patient cohort

In order to evaluate the robustness of the identified markers for metastatic disease, we measured all 743 microRNAs in an independent cohort consisting of sera from 54 neuroblastoma patients.

There is a significantly positive correlation of localized versus metastatic (stage 4) fold change abundance between samples from the first and the second cohort (Pearson’s r = 0.7, P<0.001). If only differential miRNAs between localized and metastatic disease are taken into account, the correlation between the two independent sample cohorts is even higher (Pearson’s r = 0.91, P<0.001).

In Figure 4a, a distribution of abundance fold changes is shown for localized versus metastatic disease. Seven of the 9 miRNAs identified on the first cohort have a four-fold abundance difference (2 log2 units) in the second sample cohort as well. The remaining two miRNAs show fold differences larger than 2.5 (1.4 log2 units).

**Figure 4:**
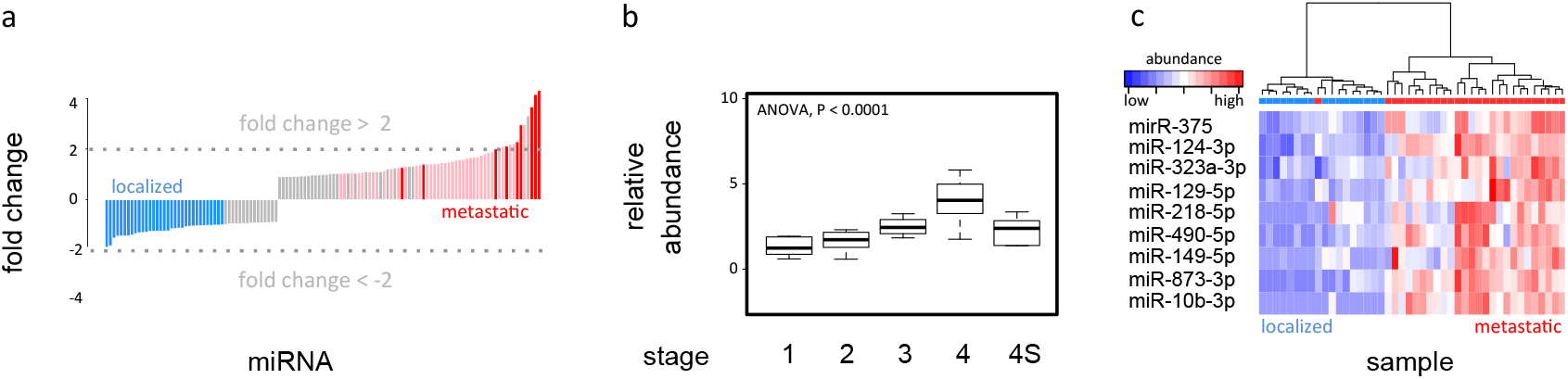
**a)** Fold change distribution of miRNA abundance between sera from metastatic (stage 4) neuroblastoma (n =27) and localized (stage 1 and 2) neuroblastoma (n=17) patients (cohort 2, n=54). Red bars indicate the position of the 9 miRNAs identified in the first cohort. **b)** Boxplot of miRNA abundance for the selected 9 miRNAs in sera from stage 1, 2, 3, 4 and 4S disease, in the cohort 2. **c)** Heatmap depicting hierarchical clustering of serum samples based on the selected set of 9 miRNAs.

A proportional increase in the average expression from the 9 miRNAs from stage 1 to stage 4 was confirmed in the second cohort (Figure 4b). An ANOVA test revealed a significant P-value of 5.8 × 10^−14^. All differences between stage 4 samples and samples from other tumor stages were significant (P-value < 0.01). In Figure 4c, the ability of the 9-miRNA set to classify the independent cohort into localized and metastatic disease is shown.

### Serum microRNA markers for metastatic disease are specific for neuroblastoma

We further evaluated the specificity of the selected set of 9 miRNAs in serum pools from healthy children and other pediatric cancer entities (sarcoma, nephroblastoma, and rhabdomyosarcoma), as depicted in Figure 5a and 5b. Low to absent levels were observed in healthy serum. When compared to serum from other pediatric cancer entities, expression levels for 8 out of 9 miRNAs were higher in neuroblastoma serum than in the pools from other pediatric cancer entities.

**Figure 5:**
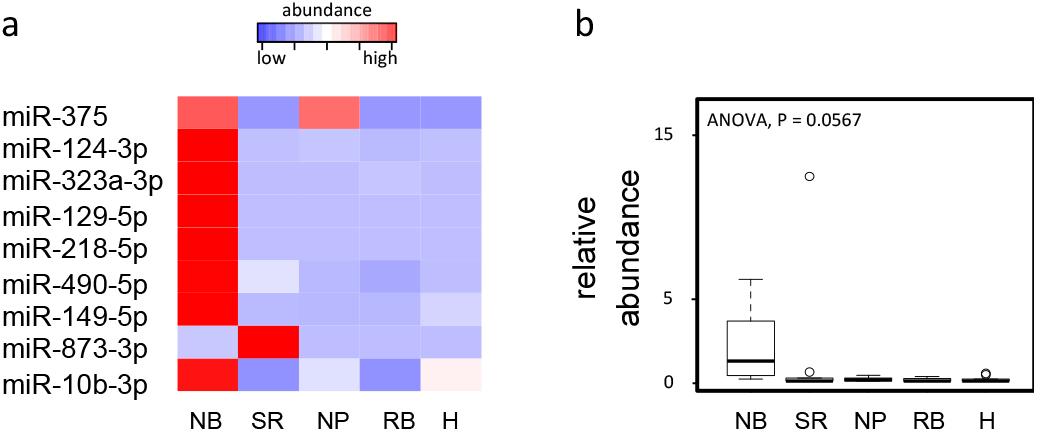
**a)** Standardized relative abundance for individual circulating miRNAs in serum pools from neuroblastoma (NB), sarcoma (SR), nephroblastoma (NP) rhabdomyosarcoma (RB) patients and healthy children (H). **b)** Relative abundance of circulating miRNAs in serum pools from neuroblastoma (NB), sarcoma (SR), nephroblastoma (NP) rhabdomyosarcoma (RB) patients and healthy children.

These results suggest that the increased levels of the identified set of miRNAs in serum from children with neuroblastoma is likely caused by the presence of neuroblastoma tumor cells in the patient’s body.

### Proportional increase of metastatic disease miRNA markers in serum from mice engrafted with human neuroblastoma cells

To evaluate to what extent tumor volume may impact the levels of miRNA markers for metastatic, we engrafted four-to-six-week-old immunodeficient mice with human luciferase-positive SH-SY5Y neuroblastoma cells. Small RNA sequencing was performed on serum samples collected 6 days before engraftment, and 11 days and 25 days after engraftment. Abundance levels of the selected set of 9 miRNAs was evaluated, in order to determine how the tumor graft impacted circulation of these miRNAs in murine serum. Significantly increased miRNA levels over time were observed for the selected miRNAs (Figure 6a and 6b). Of note, for miR-873-3p, the only miRNA that is not conserved between human and mouse, no detection was found prior to engraftment, with increasing levels 11 and 25 days post engraftment (Figure 6c).

**Figure 6:**
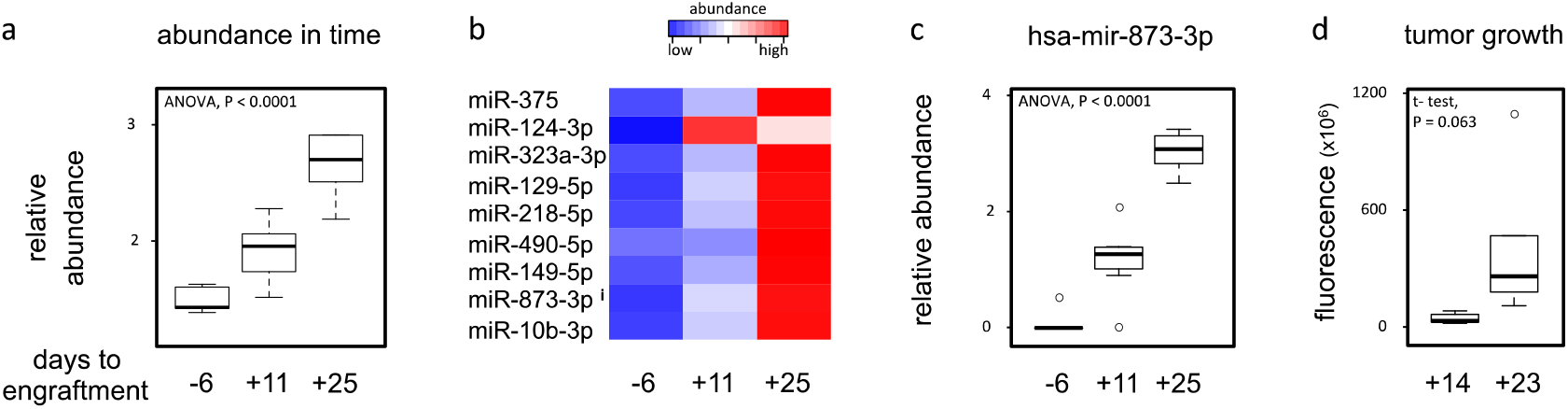
**a)** Average miRNA abundance of 9 metastatic miRNAs in murine serum from mice, 6 days before engraftment, 11 days and 25 days after engraftment with human neuroblastoma tissue. **b)** Expression of the individual miRNAs in murine serum from mice 6 days before engraftment, 11 days and 25 days after engraftment. (i) Non-conserved miRNA between mouse and human. **c)** Relative abundance of human miRNA miR-873-3p in murine serum 6 days before engraftment, 11 days and 25 days after engraftment. **d)** Luciferase fluorescence assessment of tumor volume, 14 days and 23 days after engraftment.

The higher detection levels of the selected miRNAs between 11 and 25 days post engraftment were associated with substantial tumor growth as evidenced by increased luciferase bioluminescence (Figure 6d).

These findings indicate that an increase in tumor volume results in increased abundance of the 9 miRNAs in murine serum; at least one of the miRNAs (human specific miR-873-3p) must directly originate from the growing human tumor in the immunodeficient mice.

### Serum markers for metastatic disease are high but non-differential in primary tumors

Two public datasets of miRNA expression in fresh-frozen diagnostic neuroblastoma tumors were used to evaluate expression of the serum markers for metastatic disease in the tumor. The tumor expression datasets were generated using different miRNA expression profiling technologies than used in the present study: (i) stem-loop RT-qPCR miRNA expression method (Life Technologies), as described in Mestdagh et al. (2008) and (ii) small RNA sequencing as described in Schulte et al. (2010).[8,9]

There were 201 microRNAs measured in common between our serum dataset and the stem-loop RT-qPCR tumor dataset, including 6 of the 9 serum markers for metastatic disease. These 6 miRNAs were among the top 55% miRNAs with the highest expression (Figure 7a). Fold changes were determined between 25 stage 4 tumors from patients with fatal outcome, and 25 stage 1 tumors from patients with favorable outcome. No significant correlation was found between serum and tumor microRNA fold changes (Pearson = − 0.08, P-value = 0.22) (Figure 7b). While only 3/6 miRNAs (miR-129-5p, miR-149-5p, and miR-323a3p) were significantly differentially expressed between metastatic and localized disease in the tumor dataset, it is striking that the fold changes are in opposite direction as observed in circulation: in serum, the miRNAs are more abundant in metastatic disease, in tumor they are higher expressed in localized disease.

**Figure 7:**
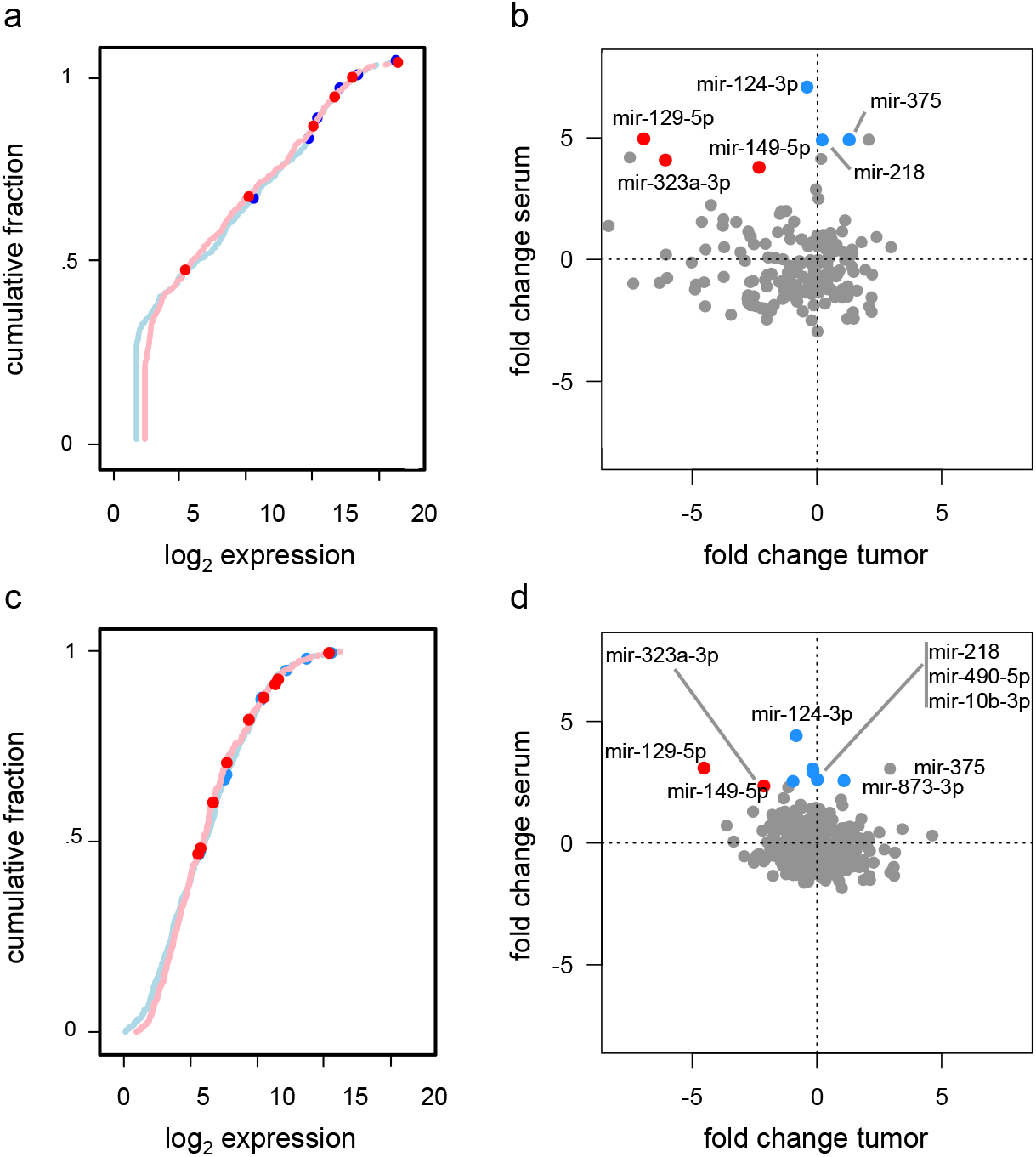
**a)** Cumulative fraction of miRNA expression in the dataset processed by stem-loop RT-qPCR tumor. Red indicates average expression in metastatic (stage 4) disease and blue indicates average expression in localized (stage 1) disease. Red and blue dots indicate the position of the serum markers for disease burden in metastatic and localized disease, respectively. **b)** Correlation of miRNA abundance differences in serum and in tumor tissue measured by stem-loop RT-qPCR. The names of the serum markers for disease burden are indicated; red dots represent differentially expressed miRNAs between metastatic and localized tumors; blue dots represent non-differentially expressed miRNAs. **c)** Cumulative fraction of miRNA expression in the small RNA sequencing tumor dataset. Red indicates average expression in metastatic disease and blue indicates average expression in localized disease. Red and blue dots indicate the position of the serum markers for disease burden. **d)** Correlation of miRNA abundance differences in serum and in tumor tissue measured by small RNA sequencing. MiRNAs in serum as markers for disease burden are indicated; red dots represent differentially expressed miRNAs between metastatic and localized tumors (P-value < 0.5); blue dots represent non-differentially expressed miRNAs.

In the small RNA sequencing dataset, there were 5 patients with metastatic (stage 4) disease and fatal outcome and 5 patients with localized disease (stage 1) and favorable outcome. All 9 serum markers for disease burden were present in the tumor dataset. These miRNAs were among the top 54% miRNAs with highest expression in the tumor (Figure 7c). No positive correlation was found between serum and tumor microRNA fold changes (Pearson = − 0.13, P-value = 0.01) (Figure 7d). Only 2/9 miRNAs (miR-129-5p and miR-149-5p) showed a significantly differential expression between metastatic and localized disease in the tumor dataset. Again, while in serum these miRNAs are more abundant in metastatic disease, in tumor they were found to be higher expressed in localized disease.

## Discussion

In this study, the abundance levels of 743 human miRNAs were evaluated in 185 diagnostic serum samples from 2 neuroblastoma patient cohorts. The association between circulating miRNA abundance variance and patient characteristics in the 1^st^ cohort revealed that tumor stage had the strongest impact on miRNA levels. By taking the overlap from the differential abundance analysis when comparing metastatic stage 4 and localized stages 1 and 2 disease and the outcome of the variance modeling, a core set of 9 miRNAs was defined. These miRNAs were highly abundant in serum from patients with metastatic disease in both tested patient cohorts. Evaluation of the selected set of 9 miRNAs in serum from stage 1,2, 3, 4 and 4S disease showed that the more the disease is spread throughout the body, the higher the levels of the selected miRNAs in serum are. This significant association to tumor stage was confirmed in both patient cohorts and we therefore decided to label this set of miRNAs as potential circulating biomarkers for metastatic disease in neuroblastoma.

To evaluate to what extent tumor load could influence circulating miRNA levels, we measured their abundance in serum from mice orthotopically engrafted with human neuroblastoma cells. A clear increase of the core set of 9 serum miRNA levels was observed 11 days and 25 days after orthotopic injection. Interestingly, there was one non-conserved (human-specific) miRNA which was only detected in murine serum after human neuroblastoma engraftment (first time point, 11 days after engraftment), with even more pronounced abundance two weeks later (25 days after engraftment). Furthermore, the increased levels of the core set of 9 miRNAs in murine serum were accompanied by substantial growth of the human tumor in the murine host. Not only do these findings show that the increased levels of the core set of miRNAs in serum is triggered by the growing neuroblastoma tumor, but they also provide clear evidence that at least one of these miRNAs is derived from the tumor tissue itself.

Comparison of neuroblastoma serum with serum from healthy children or from children with other cancer types showed that the core set of miRNAs is primarily detected at high levels in neuroblastoma serum, indicating that these miRNAs may be specific to neuroblastoma disease. In a similar study, performed by Murray et al. (2015),[6] 5 miRNAs were shown to be specifically abundant in serum from *MYCN*-amplified high-risk neuroblastoma compared to serum from non-*MYCN*-amplified low-risk neuroblastoma and 9 other pediatric cancer entities. Three of these miRNAs (miR-124-3p, miR-218-5p and miR-490-5p) were also part of the core set of miRNAs identified in the present study. These confirmatory results show that the findings in our study are robust and highlight the potential of using serum miRNAs in future clinical practice since their abundance may be reliably measured independently from sample processing methodologies and miRNA profiling technologies.

In order to assess whether the circulating biomarker miRNAs are overexpressed in metastatic tumors compared to localized neuroblastoma, we evaluated their expression in two public datasets from diagnostic primary neuroblastoma samples – one established using stem-loom RT-qPCR and the other by deep sequencing of small RNAs. The comparison of abundance differences between serum and primary tumor from metastatic versus localized disease showed a reproducible lack of correlation. There were two commonly differentially expressed miRNAs between metastatic and localized disease: miR-129-p and miR-149-5p. However, in each tumor tissue dataset these miRNAs were significantly higher expressed in localized disease compared to metastatic disease, while the inverse holds true for their circulating levels in the serum. This finding strongly suggests that the miRNA levels in serum are not upregulated due to differential expression in the tumor, but are likely proportional with tumor volume, and as such reflect disease burden and metastatic status. The fact that these miRNAs were among the most abundant miRNAs in the primary tumor further supports this.

With the present study, we provide unique first insights into the poorly understood mechanisms behind serum miRNA levels and its relation to tumor miRNA expression. In summary, we have identified 9 miRNAs and convincingly demonstrated that their elevated levels in serum are correlated to metastatic neuroblastoma disease in human samples and tumor volume in serum samples from murine xenografts. Further studies are required in order to evaluate the utility of these miRNAs as potential biomarkers for disease burden in a longitudinal disease monitoring or treatment efficacy study design.

## Materials and methods

### Patient demographics

Serum was collected from 185 neuroblastoma patients at diagnosis from 6 different clinical centers: Ghent (Belgium), Essen and Cologne (Germany), Lyon (France), Tel Aviv (Israel), and Brno (Czech Republic). Clinico-pathological properties are listed in Supplemental table 1. The serum samples were collected and processed in two independent cohorts. Cohort 1 contained 131 samples from all six centers; cohort 2 contained 54 samples from Cologne. The number of samples over different tumor stages, for each sample cohort is shown in the Table 1. Serum from healthy children (5 samples) and diagnostic serum for three additional cancer entities (sarcoma, nephroblastoma and rhabdomyosarcoma, 5 samples each) were obtained from Ghent University Hospital, Ghent, Belgium.

**Table 1:** Distribution and percentage of serum samples for tumor stage 1, 2, 3, 4, and 4s, in neuroblastoma patient cohort 1 and patient cohort 2.

| tumor stage | number of patients cohort 1 | % cohort 1 | number of patients cohort 2 | % cohort 2 |
| --- | --- | --- | --- | --- |
| stage 1 | 20 | 15% | 9 | 17% |
| stage 2 | 13 | 10% | 8 | 15% |
| stage 3 | 16 | 12% | 4 | 7% |
| stage 4 | 78 | 60% | 27 | 50% |
| stage 4s | 4 | 3% | 6 | 11% |
| total | 131 |  | 54 |  |

### Serum sample pooling

Three neuroblastoma serum pools were prepared from human diagnostic samples (Supplemental table 2), each containing 200 μl from 5 serum samples (40 μl each): pool 1 contained 5 high-risk patients that died of the disease within 36 months after diagnosis; pool 2 contained 5 high-risk surviving patients with at least 5 years follow-up time; pool 3 contained 5 low-risk surviving patients with at least 5 years follow-up time. The same approach was used for preparation of three additional pools from diagnostic serum from the other pediatric cancer entities and healthy controls.

### RNA isolation, reverse transcription and RT-qPCR reaction

The serum pools were screened for 1805 microRNAs using qPCR (miRBase release version 20), as described in Zeka et al., 2015.[10] The individual samples were screened for 754 microRNAs, including 751 human miRNAs and 3 spike-in control miRNAs: cel-miR-39-3p, miRTC (miRNA reverse transcription control, included in the RT kit) and PPC (positive PCR control, synthetic spike-in control included in the SYBR Green PCR mix). Briefly, the commercial miRNeasy serum/plasma kit (Qiagen) was used to extract RNA from 200 μl serum. The serum samples were lysed with 1000 μl Qiazol and spiked with 3.5 μl synthetic cel-miR-39-3p (1.6 × 10^8^ copies/μl). 200 μl chloroform was used for phase separation at 4 °C and 12 000 g. 600 μl of the aqueous phase was transferred into an RNeasy MiniElute spin column and subsequently washed with 100% ethanol, buffer RWT, buffer RPE and 80% ethanol. The total RNA fraction was eluted with 14 μl RNase-free water.

For reverse transcription (miScript II RT kit, Qiagen), 1.5 μl RNA was mixed with 1 μl reverse transcriptase, 2 μl RT buffer, 1 μl nucleic acid mix and 4.5 μl RNase-free water. Samples were incubated for 60 minutes at 37 °C and 5 minutes at 95 °C.

2 μl of the RT product was diluted 22-fold with 42 μl nuclease-free water and used for quantification of synthetic spike-in RNA molecules: cel-miR-39-3p and miRTC in 10 μl qPCR reactions. To this purpose, 2 μl of the diluted RT product was mixed with 5 μl SYBR Green PCR mix, 1 μl universal primer, 1 μl miRNA specific primer and 1 μl RNase-free. Samples were incubated at 95 °C for 15 minutes, followed by 40 cycles of denaturation (15 seconds at 94 °C) annealing (30 seconds at 55 °C) and extension (30 seconds at 70 °C).

For the preamplification reaction (miScript PreAMP PCR Kit, Qiagen), 5 μl of 5-fold diluted RT product was mixed with 5 μl preamp buffer, 2 μl DNA polymerase, 5 μl target specific assay pool, 1 μl universal primer mix and 7 μl RNase-free water to obtain a 25 μl preamplification reaction. Samples were incubated at 95 °C for 15 minutes, followed by two cycles of denaturation (30 seconds at 94 °C) annealing (60 seconds at 55 °C) and extension (60 seconds at 70 °C) and 10 additional cycles of denaturation (30 seconds at 94 °C) and one-step annealing/extension (3 minutes at 60 °C). 100 μl nuclease-free water was added to the 25 μl preamplification product in order to obtain a 5-fold dilution.

For miRNA quantification on the preamplified product, 5 μl qPCR reactions were prepared for each miRNA and each sample. 384-well plates, pre-spotted with 1X miRNA qPCR primer assays were used. qPCR mix was prepared for each multi-well plate by mixing 1025 μl SYBR Green PCR mix, 205 μl universal primer mix, 50 μl preamplified template cDNA, 770 μl nuclease-free water and dispensed by a Tecan Evo75 liquid handler (5 μl per well). The cycling conditions for quantitative PCR consisted of an initial activation step (15 minutes at 95 °C) and 40 cycles of denaturation (15 seconds at 94 °C), annealing (30 seconds at 55 °C) and extension (30 seconds at 70 °C). The synthetic spike-in control PPC was included in the SYBR Green PCR mix PPC.

All qPCR reactions were performed on a CFX384 Real Time PCR Detection System (Bio-Rad). Cq-values were determined by the CFX software manager version 3.1 with the regression Cq-value determination mode.

### Quality control of the RT-qPCR data

Quantification of cel-miR-39-3p, added to the serum samples during lysis, was used as processing control of the serum RNA isolation. Two samples with no detection of cel-miR-39-3p were excluded from the study. MiRTC level differences between serum samples and negative control samples (water mixed with MS2 phage RNA) were used to evaluate RT efficiency and qPCR inhibition. 22 serum samples were excluded due to a miRTC Cq-value of 1 cycle higher than the negative control samples. PPC levels across all 384-plates were used to assess deviating global Cq shifts as result of inter-run variation, batch-effect or technical failure.

The Cq detection cut-off was determined based on evaluation of single positive signals obtained from replicate miRNA expression data, as described in Mestdagh et al. (2014).[11] In the current study, two neuroblastoma sample pools were used instead of technical replicate samples. The fraction of single positive results was reduced by 95% at Cq-value 29, which was chosen as Cq detection cut-off.

### Collection and processing of murine serum samples

Expression data from murine serum was obtained from Van Goethem et al. (2017) [12]. In brief, miRNA expression profiles were generated through small RNA sequencing of serum samples from female mice bearing orthotopic neuroblastoma xenografts. 1 × 10^6^ human luciferase-SH-SY5Y neuroblastoma cells were surgically implanted in athymic immunodeficient NCr nude mice. Tumor size was assessed 14 days and 23 days after injection of neuroblastoma cells, by measurement of luciferase intensity. 100 μl blood was collected, 6 days before, 11 days after and 25 days after tumor cell injection in serum BD Vacutainer tubes. 50 μl of the resulting serum was used for RNA isolation with the miRNeasy serum/plasma kit (Qiagen). Subsequently, tRNA fragment depletion was performed as described in Van Goethem et al. (2016). [13] The tRNA depleted RNA samples were then suspended in 7.5 μl RNAse-free water and used for small RNA library preparation by TruSeq small RNA library preparation kit v2 (Illumina). Library size selection was performed on a Pippin Prep (Sage Science) device with a specified collection size range of 125-153 bp. Libraries were further purified and concentrated by precipitation and quantified using qPCR. Subsequently, equimolar library pools were prepared, and further diluted to 4 nM. The pooled library was sequenced at a final concentration of 1.2 pM on a NextSeq 500 using high output v2 kits (single-end, 75 cycles, Illumina). Quantification of small RNAs was done using Cobra, Biogazelle, Ghent).

### Preprocessing and data analysis

In order to define the neuroblastoma serum miRNome, pairwise comparisons of miRNA expression of the three serum pools was performed to select miRNAs detected in at least one of the serum pools based on three selection criteria. First, miRNAs with ΔCq larger than 1.5 between two pools were selected (303 miRNAs). Second, miRNAs showing Cq-values below 27 in one sample and an absent signal (Cq > 29) in the other sample were selected (90 miRNAs). The final selection criterion was based on detection (Cq < 28) in the high-risk serum pools (358). This resulted in a total of 751 unique miRNAs that were subsequently profiled in a first patient cohort of 131 individual serum samples. A second patient cohort containing 54 neuroblastoma serum samples was independently collected at a later stage of the study and processed by a different batch of miRNA expression profiling reagents.

A threshold Cq-value of 29 was determined based on the evaluation of single positive signals obtained from replicate miRNA expression data, as described in Mestdagh et al. (2014). [11]

After removal of Cq-values above 29 cycles, miRNAs with missing values in more than 75% (an empirically chosen threshold) of the samples were excluded from the dataset. This led to exclusion of 8 miRNAs.

Raw Cq data were normalized as described in D’haene et al., (2012). [14] Briefly, the arithmetic mean Cq for each miRNA across all samples was determined and subtracted from individual Cq-values (i.e. mean centering of the miRs). Then, the arithmetic mean Cq value per sample was calculated across all miRNAs and subtracted from each individual Cq-value to obtain normalized miRNA expression values per sample.

Generalized additive modeling, Mann-Whitney U test, Student’s T-test and Analysis of variances (ANOVA) was performed using R statistical programing tool version 3.3.2.

For the evaluation of the variance of miRNA abundance in serum by GAM analysis, the following mathematic model was used:

*miRNA expression_i_ = β_1*_INSS tumor stage + β_2*_MYCN status + β_3*_survivai + β_4*_age at diagnosis*

*For: i = miRNA 1 to miRNA 743*

*β (1 to 4) = coefficient of the prediction model for INSS tumor stage (stage 1, 2, 3, 4, 4S coded from 1 to 5), MYCN status (normal=0, amplified=1), survival (alive=0, dead=1), age at diagnosis (younger than 18 months=0, older than 18 months=1)*

### Author contributions

Concept and design of the study: Fjoralba Zeka, Pieter Mestdagh, Jo Vandesompele, Katleen De Preter. Acquisition and analysis of data: Fjoralba Zeka, Pieter Mestdagh, Jo Vandesompele. Interpretation of data: all authors. Manuscript drafting: Fjoralba Zeka, Pieter Mestdagh, Jo Vandesompele. Manuscript revision: all authors. Final approval of manuscript: all authors.

## Acknowledgements

The authors express their special gratitude to the Fournier-Majoie foundation for the generous support and guidance throughout the course of the study. The authors also thank the Flemish League Against Cancer (KOTK), Foundation against Cancer (STK), Belgian National Lottery for contributing to this work, the National Health Institute Carlos III and the European Regional Development Fund (ISCIII&FEDER).

